# Real-time Quantification of in vivo cerebrospinal fluid velocity using fMRI inflow effect

**DOI:** 10.1101/2023.08.14.553250

**Authors:** Tyler C. Diorio, Vidhya Vijayakrishnan Nair, Neal M. Patel, Lauren E. Hedges, Vitaliy L. Rayz, Yunjie Tong

## Abstract

*In vivo* estimation of cerebrospinal fluid (CSF) velocity is crucial for understanding the glymphatic system and its potential role in neurodegenerative disorders such as Alzheimer’s disease and Parkinson’s disease. Current cardiac or respiratory gated approaches, such as 4D flow MRI, cannot capture CSF movement in real time due to limited temporal resolution and in addition deteriorate in accuracy at low fluid velocities. Other techniques like real-time PC-MRI or time-spatial labeling inversion pulse are not limited by temporal averaging but have limited availability even in research settings. This study aims to quantify the inflow effect of dynamic CSF motion on functional magnetic resonance imaging (fMRI) for *in vivo*, real-time measurement of CSF flow velocity. We considered linear and nonlinear models of velocity waveforms and empirically fit them to fMRI data from a controlled flow experiment. To assess the utility of this methodology in human data, CSF flow velocities were computed from fMRI data acquired in eight healthy volunteers. Breath holding regimens were used to amplify CSF flow oscillations. Our experimental flow study revealed that CSF velocity is nonlinearly related to inflow effect-mediated signal increase and well estimated using an extension of a previous nonlinear framework. Using this relationship, we recovered velocity from *in vivo* fMRI signal, demonstrating the potential of our approach for estimating CSF flow velocity in the human brain. This novel method could serve as an alternative approach to quantifying slow flow velocities in real time, such as CSF flow in the ventricular system, thereby providing valuable insights into the glymphatic system’s function and its implications for neurological disorders.

## INTRODUCTION

The recently proposed glymphatic system involves the interchange of cerebrospinal fluid (CSF) and interstitial fluid within the perivascular space resulting in the elimination of waste from the central nervous system^11,13^. The network of perivascular spaces surrounding blood vessels in the brain facilitates the clearance of metabolic waste products, such as beta-amyloid, from the brain^10^. This system is unique in that its function is elevated during sleep, when the brain is at rest^24^. The malfunctioning of the glymphatic system has been associated with the buildup of harmful proteins in the brain, potentially resulting in a decline in cognitive function and the degeneration of nerve cells. Recent studies have shown that the glymphatic system may play a role in the development and progression of neurological disorders such as Alzheimer’s disease, Parkinson’s disease, and multiple sclerosis^2,18,32^. Researchers are actively working to bridge the gap between the coupling of the microscale glymphatic system to the macroscale CSF flow system^19–21^.

As a result, attention has been turned to understanding the flow of CSF within the brain and how it is affected by cardiac, respiratory, and low frequency oscillations^4,22^. Previous work has shown that respiration is a major regulator of CSF flow, with physiological alterations by respiratory pattern driving changes in the magnitude and directionality of CSF flow^3,15,22,27^. Hypercapnia, or an increase in the level of carbon dioxide in the blood, can be induced through respiratory patterning, such as breath holding, and causes arterial vasodilation in the brain. As per the Monro-Kellie doctrine, changes in a volume component, such as arterial vasodilation, within the fixed volume of the skull causes compensatory changes in at least one other components, such as CSF outflow^16^.

MRI flow measurements are often utilized as a non-invasive approach to visualize and quantify the macroscale flow of CSF, which can be used to identify alterations across diverse patients and pathologies. MRI flow imaging sequences, such as planar or 3-directional phase-contrast MRI (4D Flow MRI), can be used to detect CSF motion but are restricted within one cardiac/respiratory cycle due to gating constraints^28^. Only a few studies observe real-time CSF movement, using either real-time phase-contrast techniques^23,29^ or time-spatial labeling inversion pulse^25,26^. Additionally, these sequences are often not available at research scanners since they are uncommon or new imaging sequences. Another imaging technique, blood oxygen level-dependent functional magnetic resonance imaging (BOLD fMRI) is sensitive to changes in the magnetic properties of blood as it flows through the brain^9^. In the context of CSF flow, BOLD fMRI can be used indirectly to detect real-time changes in the velocity of CSF flow through the brain ventricles and subarachnoid spaces by accounting for inflow effect-mediated signal increases.

Fultz et al. utilized BOLD fMRI (TR < 400 ms), to study the temporal patterning of electrophysiology, hemodynamics, and CSF movements^4^. Their study maximized inflow effect mediated signal increases by carefully placing the first slice of their imaging volume at the bottom of the fourth ventricle, where cranial-directed flow-mediated inflow effect would be greatest. Their work described a critical velocity value that is required to completely replace the CSF in a given voxel, however they did not delve into quantifying the underlying velocity.

Using a combination of flow physics, MR imaging experiments, and human CSF fMRI data, our present study seeks to further understand and quantify the inflow effect of dynamic CSF motion on fMRI for *in vivo* measurement of real-time CSF flow velocity. Specifically, we assessed linear and nonlinear models of velocity predictions by empirically fitting to fMRI data from a controlled flow experiment. To assess the utility of this methodology in human data, we also computed velocities on CSF flow data from eight healthy volunteers, where breathing protocols were used to amplify CSF flow oscillations.

## METHODS

### The inflow effect on fMRI

The inflow effect refers to the increase in MRI signal intensity that occurs when fresh fluid (not exposed to radiofrequency pulses) enters a region of the imaging volume, as demonstrated in **Figure 1** for various inflow scenarios. Each of the scenarios in **Figure 1** corresponds to a specific increase of the signal, which we seek to quantify. As fresh fluid moves into the imaging volume between RF pulses, it induces higher signal intensity via apparent T1 shortening. The inflow effect is distinct from the BOLD effect in origin, and its contribution to the fMRI signal can distort the hemodynamic response estimated in an event-related fMRI experiment^7^. The inflow effect is more significant when a short repetition time (TR) and large flip angle are used in a single-shot gradient-echo echo-planar imaging (EPI) sequences. The inflow contribution to the fMRI signal increases monotonically in spoiled gradient-echo sequences, while the fMRI signal may increase or decrease in conventional refocused gradient-echo sequences^7^.

**Figure 1:**
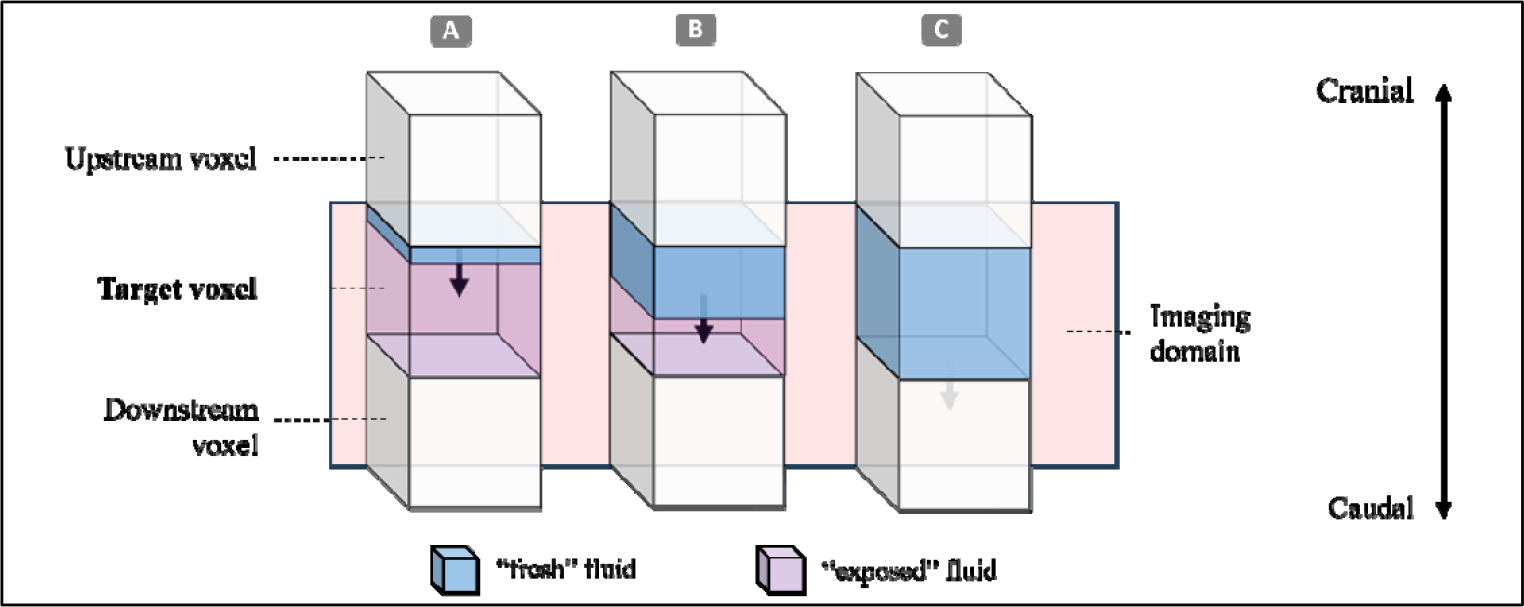
Graphical representation of possible configurations of flow changes between temporal fMRI acquisitions, where blue shading represents fluid that was originally outside of the imaging domain (termed “fresh”), and purple represents fluid that was inside the imaging domain (termed “exposed”) at the previous time step. The three stacks (A-C) represent minimal (A), majority (B), and full (C) amounts of “fresh” fluid flowing caudally from the upstream voxel.

Given that each voxel holds a volume defined by the spatial resolution of the imaging domain, we theorize that the maximum possible signal increase occurs when the full volume of a given voxel has been “refreshed” with incoming fluid in each repetition time (TR), shown as Stack C in **Figure 1**. This would be accomplished by a voxel-average critical fluid velocity (v_crit_), which is equal to the edge length (z) perpendicular to the fluid flow direction divided by the repetition time for an fMRI EPI scan:

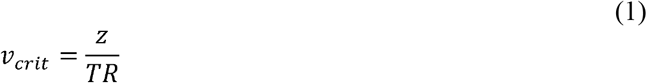

Assuming that v_crit_ corresponds to a maximum signal observed on fMRI, we can use a simple linear theoretical model to estimate instantaneous velocity, v(t) by scaling the normalized instantaneous fMRI signal (f(t)) by the critical velocity, as demonstrated below:

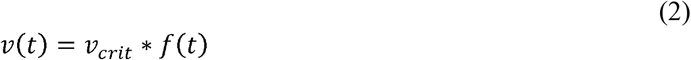

Further, previous work done by Gao et al. 1988 has demonstrated that this relationship becomes nonlinear at flip angles less than 90°, demonstrated graphically from a replication of Gao 1988 (see **Supplemental Figure S1**). The relationship, herein referred to as the nonlinear model, becomes the following when rearranging to solve for v(t) under the assumption of a plug flow spatial velocity profile (see **Appendix 1** for mathematical derivations):

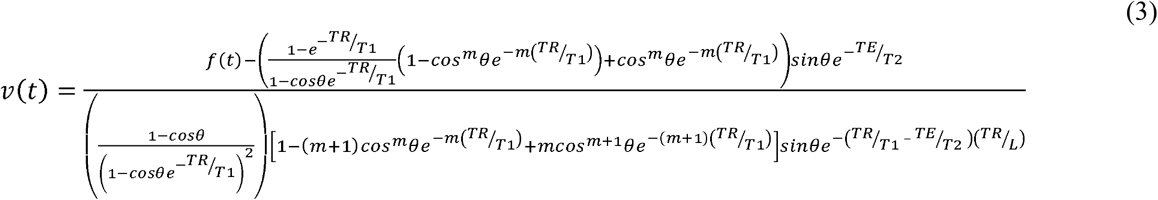

Where *TR* is repetition time, *TE* is echo time, *T1* is the spin lattice relaxation time of CSF, *T2* is the effective transverse relaxation time of CSF, θ is the flip angle, *L* is the slice thickness, and *m(t)* is the integer value of number of radiofrequency pulses a given volume of fluid will experience within a voxel defined by Equation 4 below.

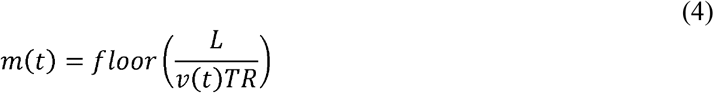

Importantly, the normalized signal *f(t)* in **Equation 3** above must match the range predicted by the nonlinear model for a given flip angle, rather than a range of zero to one. More information regarding the computation of Equation 3-4 and other aspects of the nonlinear framework are available in **Appendix 1**.

### Experimental flow study and fMRI

To test this linear model, we designed a controlled flow imaging study for a fixed set of parameters representative of the *in vivo* imaging conditions. Water was used as the working fluid for its ease of use, similarity to CSF, and conserved relaxation rates. The flow loop, partially diagrammed in Figure 2A-B, involved a programmable syringe flow pump (Legato 210, KD Scientific) loaded with a 100mL syringe fitted via Luer lock to a 4-mm inner diameter length of tubing, connected to a 10-mm inner diameter tube which was passed through the MRI Scanner (GE 3-T) and a 64-channel head coil then submerged into a fluid reservoir outlet.

**Figure 2:**
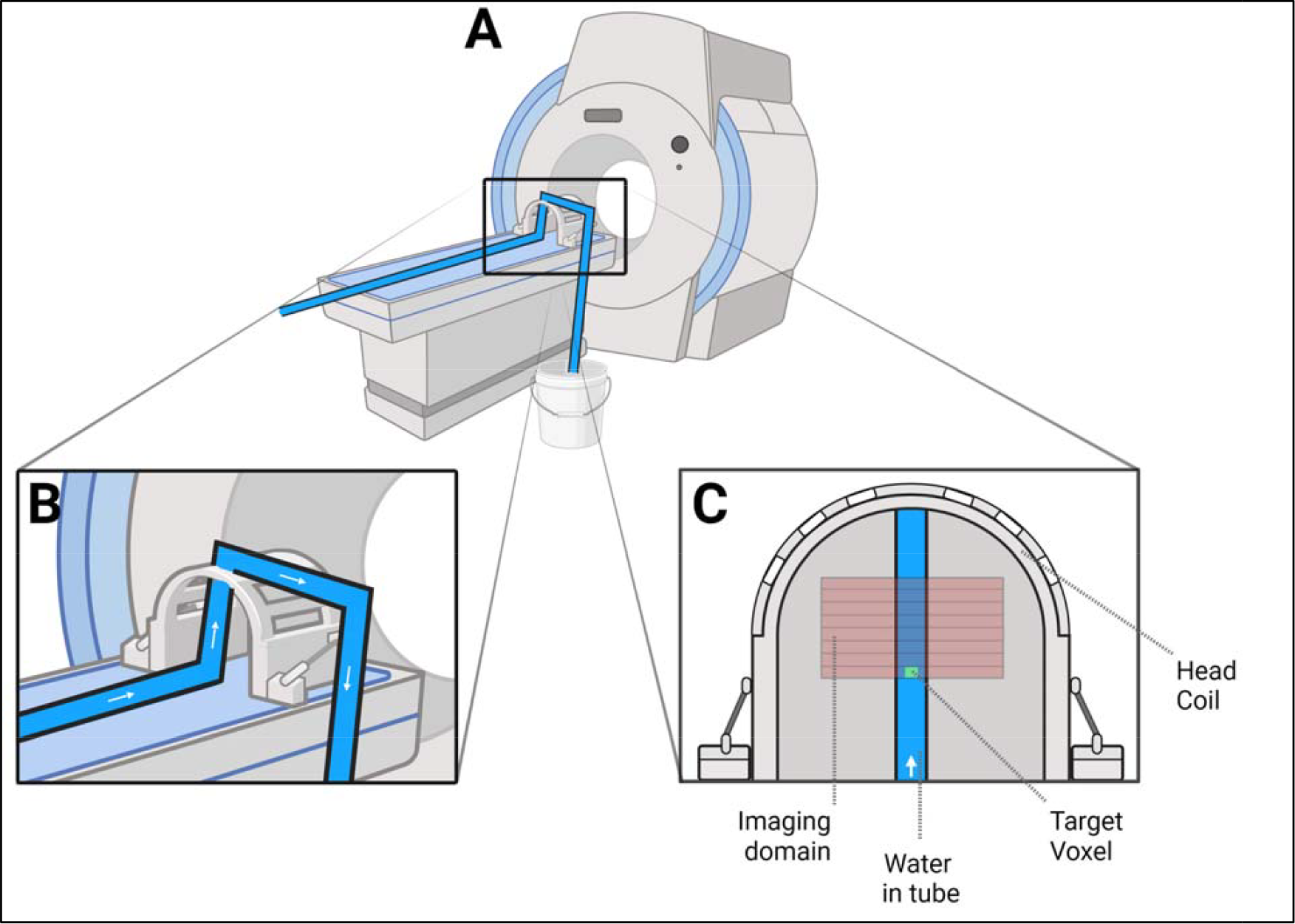
**(A)** Graphical representation of experimental flow loop and MRI scanning room set-up. **(B)** Expanded view of the flow and tube orientation within the 64-channel head coil, where blue fluid represents water and white arrow denotes steady flow direction. **(C)** Diagram view of the MRI slice selection within the head coil where the target voxel (denoted by a green square) is placed at the bottom of the imaging volume. Image created in BioRender.

The fMRI scans were acquired by using a multiband EPI sequence (FOV = 120 mm, acquisition matrix = 64 x 64, 10 slices, voxel size = 2.9 × 2.9 × 2.9mm3, TR/TE = 440/20.3 ms, echo-spacing = 0.644ms, flip angle = 35°, hyperband acceleration factor = 8, multi-slice mode: interleaved). The imaging volume was placed perpendicular to the flow direction in the head coil (**Figure 2C**), with the target voxels set on the first slice of voxels exposed to flow in the head coil, where inflow effect-mediated signal increase is greatest.

The v_crit_ for these experimental flow studies, given the isotropic spatial resolution of 2.9 mm and TR of 440 ms, was computed to be 6.59 mm/s. Thus, ten equally spaced inflow velocities, spanning 1 mm/s to 10 mm/s, were repeated over 3 trials to provide a range of velocities above and below the critical velocity within the 10-mm tube. Given the fixed 100mL volume of the syringe, rest intervals between velocity conditions allowed us to reset the syringe pump and fluid volumes.

### Experimental data processing

Measured fMRI signals for each of the three trials were analyzed by first selecting four target voxels which are aligned perpendicular to flow and in the same z-slice of the imaging volume (**Figure 3A**) to obtain a better spatial average of the flow temporal waveform. The fMRI signal was normalized for each voxel by subtracting 10^th^ percentile minimum lower bounds from the raw fMRI signal, dividing by the 95^th^ percentile maximum upper bounds, and thresholding from 0 to 1. Then, the normalized target voxels within the core of the tube were spatially averaged for each trial to better approximate the mean velocity, with standard deviation quantified across target voxels within trial (**Figure 3B**).

**Figure 3:**
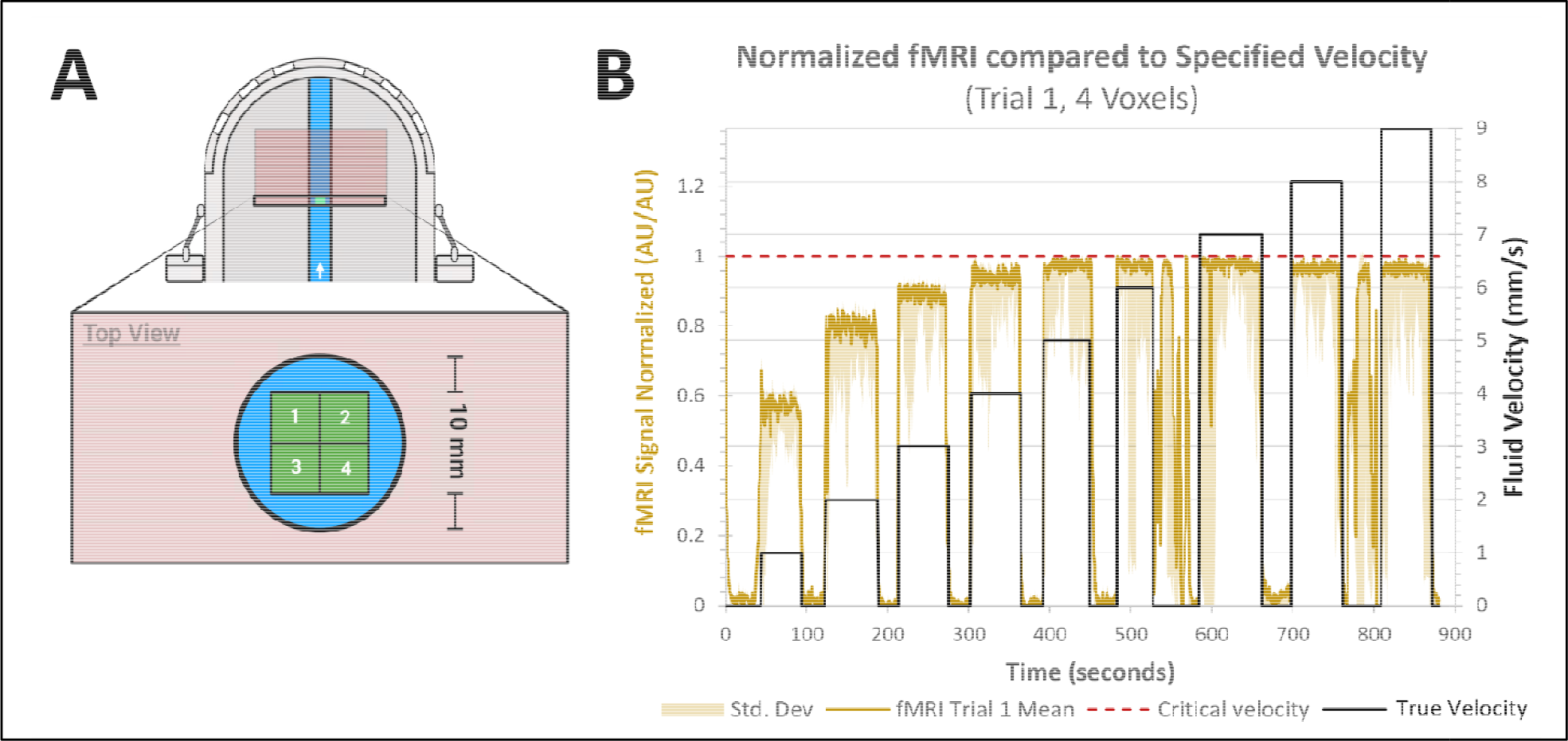
**(A)** Diagram of spatial averaging across 4 voxels center to the flow within the first slice of the imaging volume oriented perpendicular to flow. **(B)** Data comparison of temporal profiles at 1 between estimated fluid velocity (green) as set via the syringe pump and measured, normalized fMRI signal (black), where the red dashed line denotes the critical velocity at which is the target voxel would be fully refreshed with new fluid in each TR. Signal oscillations occurring between 500-600 second time points are a result of exchanging syringe volumes and resetting the syringe pump due to volume considerations.

Linear fitting was constructed using the *‘fit()’* function in MATLAB (MathWorks, R2021a) by specifying the mean measured fMRI signal, f(t), as x-data and the specified velocity, v(t), as the y-data (**Figure 4A**). A single constant linear model was specified with the constant is defined to be the critical velocity, as calculated according to the equation (2). Velocity estimations based on the nonlinear model (**Figure 4A**) were computed using Equations 3-4 above, where the value of m was informed by the known experimental fluid velocity at each flow regime in the syringe flow setup.

**Figure 4:**
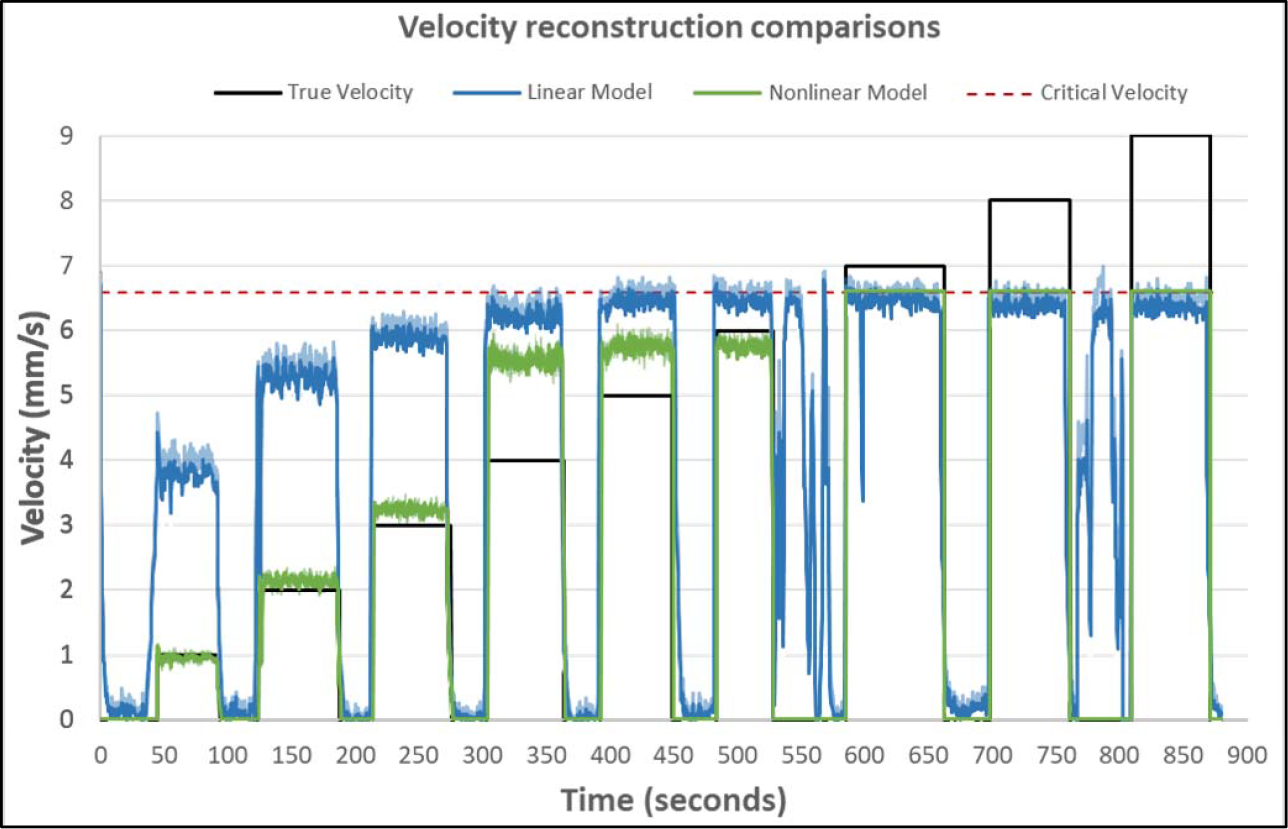
Using the linear (blue) and nonlinear (green) fits, experimental velocity profiles were reconstructed to compare against the experimental velocity data (black). Critical velocity is shown as a dashed red line.

### In vivo structural and functional MRI scans

To test the nonlinear relation of fMRI signal to velocity using the inflow effect, we recruited healthy volunteers (n=8) aged 19 – 48 (25.75 ± 9.53) years to undergo a series of structural and fMRI scans that included both resting state and breath holding regimens. Structural T1-weighted MPRAGE (Magnetization Prepared Rapid Acquisition Gradient Echo - TR/TE = 2300/2.26 ms, 192 slices per slab, flip angle = 8°, resolution =1.0mm × 1.0mm × 1.0mm) images were acquired first to accurately locate the fourth ventricle, followed by the functional scans. For the resting state scans, the volunteers were instructed via visual cues to breathe normally. For the breath holding scans, the volunteers were instructed to breathe normally (15 seconds), followed by paced breathing (18 seconds - three repeats of a 3-second inhale and 3-second exhale), and then breath hold (20 seconds) at for a total of 6 cycles. The fMRI scans were acquired by using a multiband EPI sequence (FOV = 230 mm, acquisition matrix = 92 × 92, 48 slices, voxel size = 2.5 × 2.5 × 2.5 mm3, TR/TE = 440/30.6 ms, echo-spacing = 0.51 ms, flip angle = 35°, hyperband acceleration factor = 8, multi-slice mode: interleaved) on a 3T SIEMENS Scanner at Purdue MRI Facility. The imaging volumes were placed perpendicular to the fourth ventricle with the edge of the volume at the bottom of the fourth ventricle and extending caudal into the neck as described Nair et al. 2022. Target voxels were set on the upper most (cranial) slice, where inflow effect-mediated signal increase is greatest, to understand outflow from the brain to the neck in the ventricular system.

### In vivo data processing

fMRI scans were preprocessed using FSL [FMRIB Expert Analysis Tool, v6.01; Oxford University, United Kingdom]^12^ and MATLAB (MATLAB 2020b; The MathWorks Inc., Natick, MA, 2000). As described previously^4,22,28^, preprocessing steps only included slice timing correction and registration with the structural T1-Weighted image, before extracting the fMRI inflow signal from a target voxel at the center of the fourth ventricle. Measured fMRI signals for each of the resting state and breath holding scans were analyzed by first applying detrending on the target voxel within each subject to match protocols from human data and address signal drift using the MATLAB *`detrend()`* function. The detrended fMRI signal was normalized for each voxel by subtracting 10^th^ percentile minimum lower bounds from the raw fMRI signal, dividing by the 95^th^ percentile maximum upper bounds, and thresholding from 0 to 1. The upper and lower bounds were searched within subjects, due to inter-subject variability, and across breathing patterns, as large CSF flow velocities can occur either during resting state or breath holding. For the simple linear fit, the normalized signals were scaled according to the critical velocity as described in **Equation 2**. For the nonlinear model fit, several additional pre-processing steps were required. Firstly, the normalized signals were re-scaled via linear mapping to the range predicted by the nonlinear model (visualized in Supplemental **Figure S1**). For the parameters used in the present study, this range spans from 0.2315 to 0.5626, where the lower and upper bounds refer to the signal predictions at zero and v_crit_ velocities, respectively. Given the cyclical definition of *v(t), m(t)*, and *f(t)* in the nonlinear model, where expected velocities are used to compute *m(t)* which is used to estimate signal, we sought to evaluate an empirical relationship of *m(t)* as a function of normalized signal for the range of velocities expected in this data. This was accomplished by fitting a 4-parameter exponential model to a range of 10,000 points of normalized signal predictions of known velocity values spanning from 0.1 mm/s to v_crit_ and measured signal intensities using the ‘fit()’ function in MATLAB, which returned an R^2^ value of 0.9899 for the model given by **Equation 5**. These values of *m(t)* were then used to inform the computation of **Equation 3** to compute expected velocities across the normalized, rescaled signals across all scans within each patient. More information on these calculations can be found in **Appendix 1**. In short, **Equation 5** bypasses the requirement to know the underlying velocity by fitting to the expected signal range for the two points of known velocity in our studies, when velocity is zero and when velocity is greater than or equal to v_crit_ which correspond to the minimum and maximum signal intensities respectively.

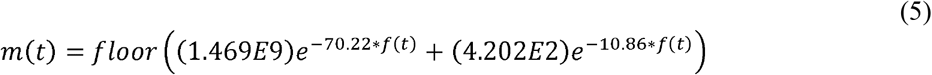

## RESULTS

Using an experimental flow loop and multiband EPI fMRI sequence, we quantified the relationship between measured fMRI signal increase due to the inflow effect and estimates of the underlying velocity of fluid, as shown in **Figure 3B**. The measured fMRI signal increase due to inflow effect related nonlinearly to the underlying velocity and notably, the nonlinear framework was found to reasonably estimate the underlying velocity in our experimental data (**Figure 4)**.

To qualitatively understand the impact of differences in recovering velocities using the linear and nonlinear models, we calculated the temporal velocity waveform using the measured fMRI signals (**Figure 4**). It can be observed that the linear model is insufficient for reconstructing velocities below the critical velocity, whereas the nonlinear model provides a closer approximation at all levels below the critical velocity for the given flip angle. It is important to note that neither model can resolve velocities above the critical velocity due to the notion that signal is proportional to unexposed fluid which is maximally refreshed at flow speeds equal to or above the critical velocity.

The linear and nonlinear models were also applied to the *in vivo* data to visually demonstrate their ability to estimate underlying velocities, from which a representative subject case as well as a group average (n=8) as shown in **Figure 5**. The nonlinear model reduces the impact of lower fMRI signal increases on reconstructed velocities as well as sharpens the temporal changes in velocity, as shown in **Figure 5C**. Both reconstructed velocity waveforms span a range from zero to the critical velocity of the *in vivo* scans. Notably, the signal and velocity waveforms follow a distinctly cyclical pattern in the breath holding data demonstrating the sensitivity of these measurements to breathing patterns known to alter CSF motion.

**Figure 5:**
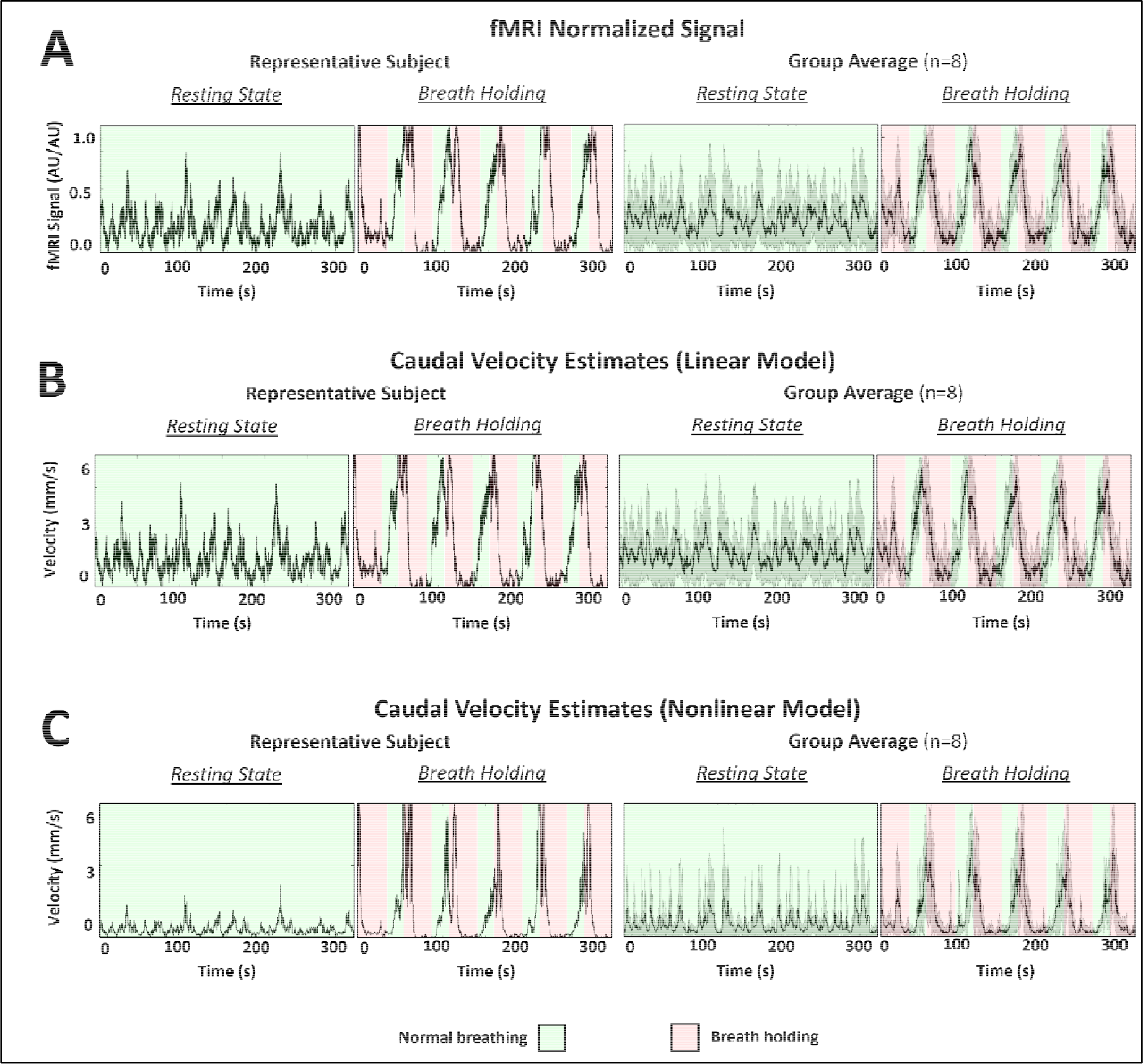
In vivo velocity temporal waveforms in a single healthy volunteer and group average (n=8), estimated from **(A)** normalized fMRI data using **(B)** linear and **(C)** Gao 1988 nonlinear fitting models. Background shading indicates Resting State free breathing (Green), where subjects were instructed to breathe normally, and Breath Holding (Red), where subjects were instructed to hold their breath.

## DISCUSSION

CSF inflow fluctuation captured by a fast EPI sequence has been discussed in detail in Fultz et al.^4^ and Yang et al.^28^, but these studies did not connect the measured inflow effect-mediated signal increases to the underlying velocities. Accurate quantification of real-time, in vivo velocity temporal waveforms could empower deeper understandings of macroscale CSF circulation and enable new metrics to stratify physiological changes in diverse patient sets. The objective of the present study was to quantify the underlying velocities that mediate the signal increase by the inflow effect for a given set of imaging parameters relevant for *in vivo* human studies of breathing regimens. This was accomplished by performing fMRI to measure inflow effect-mediated signal increase in a controlled, experimental flow loop and using the data to fit a linear and nonlinear model for reconstructing velocity waveform from fMRI sequences of similar parameters. The methods developed here provide utility for approximating slow flow velocity, e.g. ventricular CSF flow, where phase-contrast methods often perform poorly due to their restriction by a priori selection of the velocity encoding parameter. Phase-contrast methods require selection of a low velocity encoding parameter to achieve high velocity to noise ratios, but these low parameter values require higher gradients and thus can induce more noise from imaging artifacts like eddy currents.

From the data shown in **Figure 3** we observed a nonlinear increase in measured fMRI signal as a function of the underlying fluid velocity. This nonlinear behavior aligns with the behavior demonstrated in the 1988-2012 series of publications on quantifying signal increase by the inflow effect by Gao et al.^5–8^. Additionally, our reconstructions of this data quantitatively demonstrated the shortcomings of a linear model and the potential for a nonlinear model based on the Gao 1988 methodology to estimate underlying fluid velocity (**Figure 4**). Notably, both models fail to resolve any velocities greater than the v_crit_ since these velocities correspond to maximum volume replacement of the target voxel by unexposed fluid. Thus, the v_crit_ value is an upper limit for reconstructions utilizing this method. Our reconstruction method is best suited for slow moving fluid flow in relatively straight geometries due to the limitation of only capturing flow that is perpendicular to the imaging plane. The nonlinear framework utilizes a plug flow assumption, which may not generally hold true for all geometries, however in the present study voxels were spatially averaged within the region of interest^8^. This means that we are estimating a mean velocity, assumed to be uniform over the given voxels which is logically much closer approximated by a plug flow assumption rather than a spatially resolved parabolic flow assumption. If the desired flow geometry was sufficiently large and well resolved spatially, future studies could replicate the methods used in this paper using a parabolic flow assumption from Gao 1988 to reconstruct a parabolic flow profile as well.

We tested our reconstruction methods on *in vivo* human data with the region of interest in the fourth ventricle. This location is a connection for CSF transport between the midbrain and neck, relatively straight geometry which aided in perpendicular slice placement, and has broad width which minimizes partial volume effects on the target voxels of interest. A breathing regimen that included breath holding was selected to make use of the hypercapnic response of the brain, which causes increased caudal CSF outflow due to the increased volume occupied by blood during vessel dilation. Thus, the breath holding regimen was intended to boost the underlying CSF velocity above 5.68 mm/s, which is assumed to be the critical velocity of the *in vivo* imaging sequences. It is a core assumption of this model that the fluid being measured reaches the critical velocity at some point during the series of acquisitions to ensure scaling is accomplished across the correct range.

To compare the ranges of velocities calculated by the present study, it is useful to consider recent publications that measure CSF flow metrics using real-time phase-contrast MRI. Baselli et al. observed mean caudal CSF flows of 0.11 ± 0.134 mL/s^1^ at the 1^st^ cervical layer. In a more comprehensive analysis of directional and respiratory effects on CSF flow^1^, Yildiz et al. observed peak CSF flows at resting state of 1.45 ± 0.594 mL/s and in deep breathing of 2.04 ± 0.794 mL/s caudally at the foramen magnum^30^. Given their differing anatomical measurement locations, these flow rates can be scaled via conservation of mass to the level of the fourth ventricle and converted to average velocity for comparison to the values observed in the present study (see **Appendix 2)**. This results in 4^th^ ventricle velocity estimates from Baselli et al. of 0.393 ± 0.502 mm/s mean caudally and from Yildiz et al. in resting state of 5.13 ± 3.02 mm/s peak caudally and in deep breathing of 7.20 ± 4.14 mm/s peak caudally. The current study estimated group-averaged mean CSF flow velocity during resting state as 0.518 ± 0.646 mm/s and during breath holding as 0.918 ± 0.850 mm/s caudally at the fourth ventricle. These results are on par with current literature estimates on a group-wise basis. Notably, the large temporal oscillations in velocities vary across an order of magnitude which contributes to the large standard deviations. The method established in the present study represents an alternative approach to measuring CSF flow in real time.

The results of the present study may have limited generalizability outside of the imaging parameters used within the *in vivo* studies. As demonstrated in Gao 1988, the increase in inflow-mediated signal increase depends on several factors, including the pulse sequence, imaging parameters, number and position of the slices, and fluid T1/T2, which is field-strength dependent^7,8^. The current study did not evaluate changes in these parameters, which could be used to acquire a more robust reconstruction, thus our results may only be valid for our specific combination of TR (0.44 s), z (2.5 – 2.9 mm), and flip angle (35°). Further, the performance of the linear model may increase with increasing flip angles due to the increase in linearity of the nonlinear frameworks at flip angles approaching 90° (see Supplemental **Figure S1**). Although water was used as a surrogate for CSF, it is important to note that water only approximates the solute properties and T1/T2 signal of CSF. Although the experimental data was obtained on a GE 3T scanner, the human subject data was acquired on a Siemens 3T scanner, which could introduce interscanner variability. Despite these limitations, this method represents a novel alternative approach to estimating slow moving fluid velocities directly from fMRI with high temporal resolution and the absence of cardiac or respiratory gating requirements, which has the potential to enable better quantification of macroscale CSF flow in relevant patient sets for understanding neurodegenerative diseases.

## List of Abbreviations

BOLD fMRI: Blood oxygen level-dependent functional magnetic resonance imaging
CSF: Cerebrospinal Fluid
EPI: Echo-planar imaging
v_crit_: critical fluid velocity

## ACKNOWLEDGEMENTS

This study was funded by the NIA 1R21AG068962-01A1 award. The Life Science MRI Facility was funded by an NIH S10 instrumentation grant: S10 OD012336. We are grateful for the technical assistance in operation of the syringe pump from Hui Ma of Purdue University, Weldon School of Biomedical Engineering.

## Appendix 1: Gao 1988 Derivations & Nonlinear Framework

The Gao 1988 publication referred to in our work set forth two derivations for measured MRI signal on a spoiled FLASH sequence using (A) plug, and (B) parabolic flow profile assumption. The plug flow model was chosen in our study due to its innate representation of averaged data and due to our inability to assume an underlying parabolic flow profile given the complex geometry of the fourth ventricle where measurements are taken *in vivo*. Thus, equation 29 of the Gao 1988 framework, rewritten below, was used as the basis for the nonlinear model used in this work:

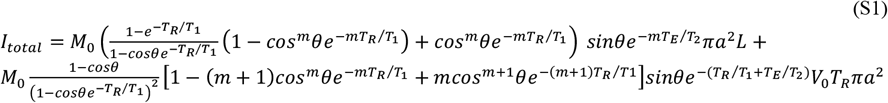

Where *I*_*total*_ is the MRI signal, *M*_0_ is the equilibrium magnetization, *T*_*R*_ is repetition time, *T*_*E*_ is echo time, *T*_*1*_ is the spin lattice relaxation time of CSF, *T*_*2*_ is the effective transverse relaxation time of CSF, *θ* is the flip angle, *V*_0_ is the average velocity, *a* is the diameter of the channel, *L* is the slice thickness, and *m* is the integer value of number of radiofrequency pulses a given volume of fluid will experience within a voxel.

Importantly, this equation can be normalized by dividing *I*_*total*_ by *M*_0_*πa^2^L* as demonstrated in Figures 4-5 of Gao et al. 1988. Rearranging Equation S1 above results in the following:

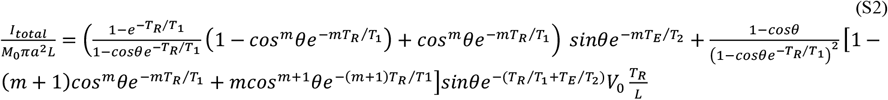

However, the range of this normalized equation depends strongly on the flip angle of the underlying MRI sequence used. Thus, in the case of the present study with a flip angle of 35° the proper normalized range varies from [0.22, 0.55] which was accomplished by mapping the [0, 1] normalized range to the nonlinear predicted range.

**Figure S1:**
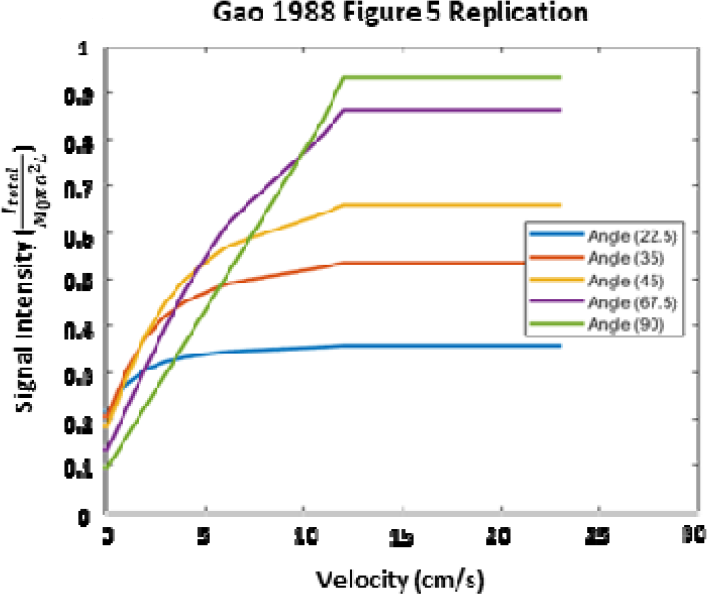
Normalized signal intensity represented as 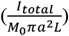 replicated from Gao et al. 1988 using an additional flip angle of 35° as well as the imaging parameters from Gao et al. 1988.

Once the fMRI data has been mapped to the proper normalized range, we can represent the nonlinear estimation of underlying velocity profile by rearranging Equation S2:

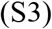

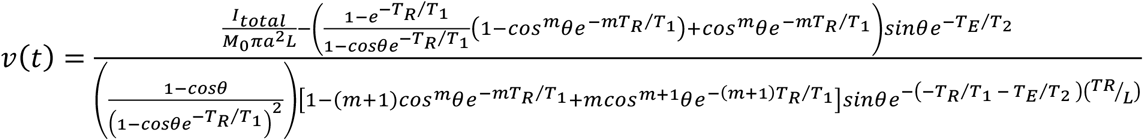

Although all imaging parameters are known and the 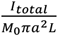 is accounted for with measured data normalized mapped, the *m* parameter remains undetermined and is described by the following Equation 22 from Gao 1988:

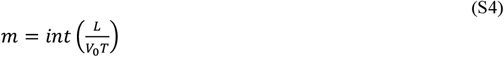

This parameter introduces a circular logic between Supplementary Equations 3-4, where underlying velocity estimates are required to reconstruct the temporal velocity profile. Thus, we sought to evaluate the relationship between *m* and 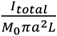 for our *in vivo* fMRI scans using the notion that the underlying velocity is known when the fluid has zero velocity and v_crit_, which correspond to minimum and maximum normalized fMRI signals respectively. Therefore, we applied the following logic:

1. Prepare a velocity range from [0.1 mm/s, v_crit_] across 10,000 data points

**Figure S2:**
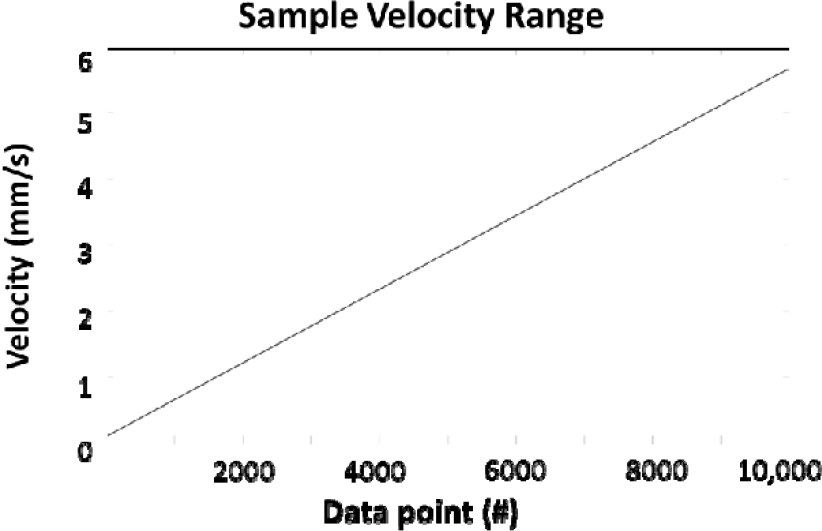
Sample velocity range spanning 0.1 mm/s to v_crit_ = 5.67 mm/s across 10,000 data points.
2. Compute nonlinear normalized and mapped signal range using scan parameters and velocity range from **1**. Using Supplementary Equation 2

**Figure S3:**
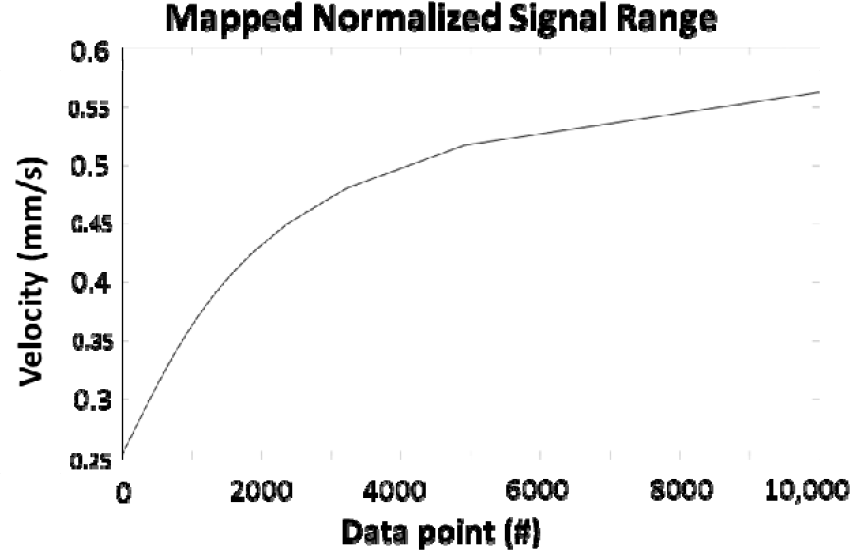
Nonlinear normalized signal range 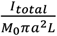 computed using Supplementary Equation 2.
3. Compute *m* at every test point then fit double exponential model to entire normalized signal range

**Figure S4:**
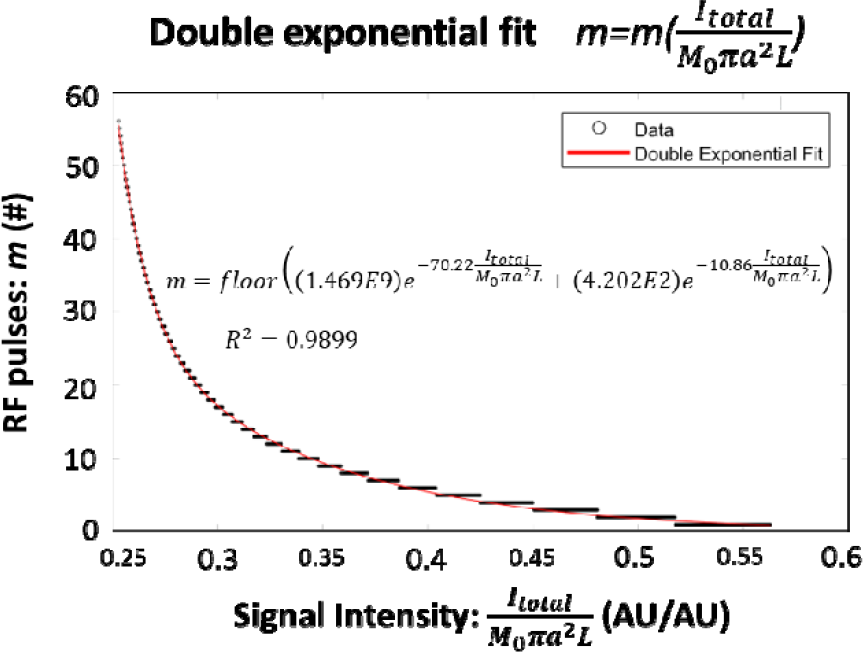
Double exponential fit of m, representing the # of RF pulses experienced by a plug of fluid flow within a given voxel, against the nonlinear normalized signal range. 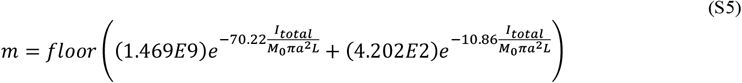
4. Compute temporally resolved nonlinear normalized signal range using scan parameters and velocity range from **1**. by the equation below:

**Figure S5:**
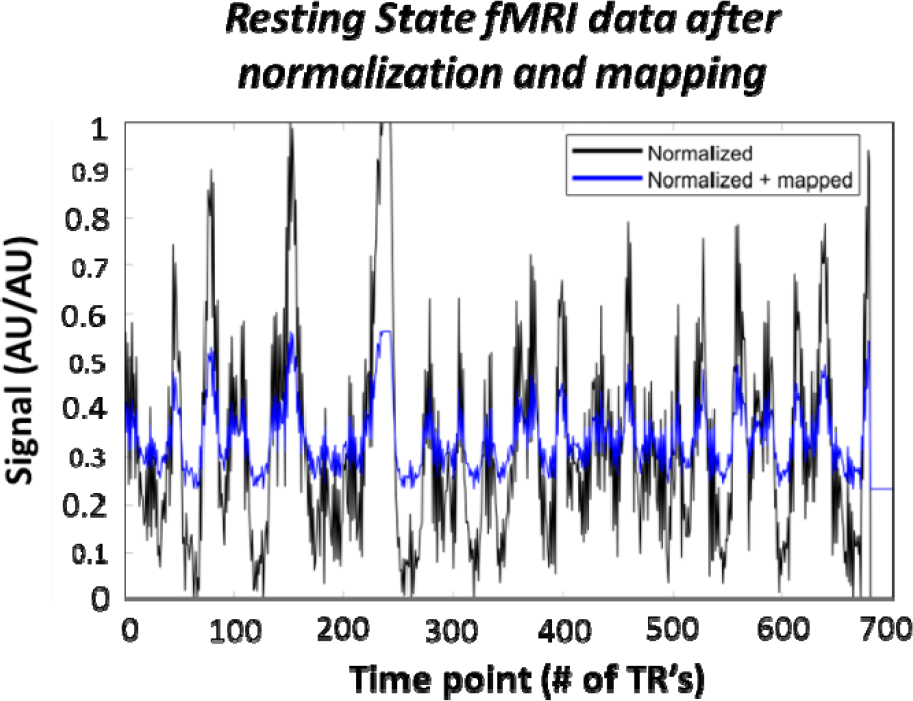
Visualization of the range of normalized data before and after mapping to the nonlinear range computed for our MRI parameters as visible in Figure 7.

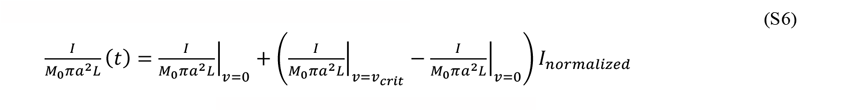
5. Use the double exponential model from **Supplementary Equation 5** alongside **Supplementary Equation 3** to recover the underlying velocity from a sample subject in our *in vivo* data

**Figure S6:**
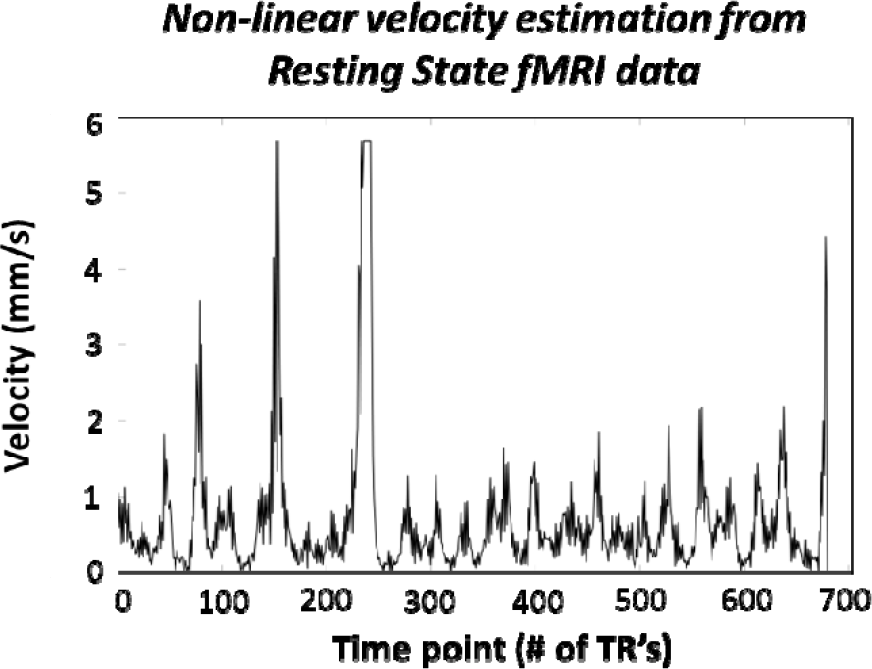
In vivo velocity temporal waveform estimated using the double-exponential fit for # of RF pulses in tandem with Supplementary Equation 5.

## Appendix 2: CSF flow and velocity estimations

Conservation of mass was used to estimate fourth ventricle fluid velocities at resting state and during breathing protocols by scaling literature values measured at other connected CSF locations. Baselli et al. measured flowrate at the 1^st^ cervical layer whereas Yildiz et al. measured velocity at the foramen magnum, both caudal to the fourth ventricle. See **Supplemental Table 1** and **Figure 11** for more details.

**Supplemental Table 1:**
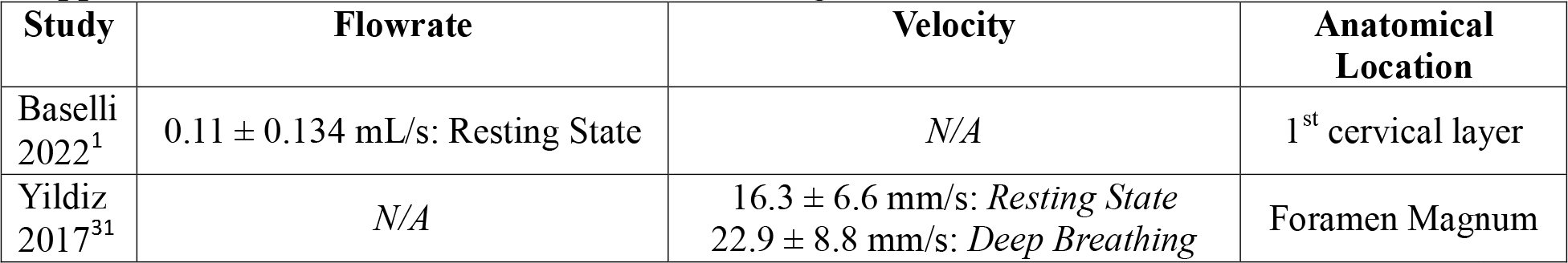
Literature values for measuring real-time CSF flow metrics

**Figure S7:**
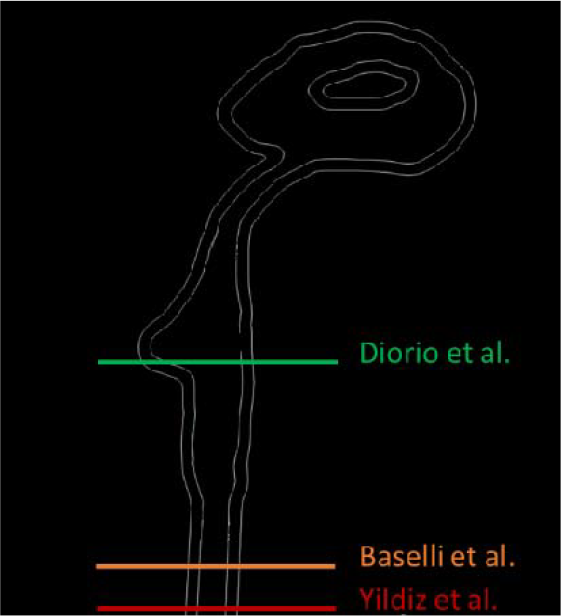
Anatomical scanning locations within the cerebrospinal fluid-filled ventricular system for 2 literature studies (Baselli et al. 2022 and Yildiz et al. 2017) compared to the current study.

By assuming negligible fluid outlet between anatomical scan locations at the fourth ventricle, 1^st^ cervical layer, and foramen magnum flow rates would be equivalent at each cross section. Thus, the Baselli et al. CSF flowrate^1^ is already provided, whereas the Yildiz et al. data requires multiplication of mean velocity by cross-sectional area at the foramen magnum, see **Supplemental Table 2**.

**Supplemental Table 2:** Literature values for ventricular system anatomical dimensions

| Study | Area | Anatomical Location |
| --- | --- | --- |
| Piaggio et al. 2018<sup>17</sup> | $88.9 \pm 6.0 \text{ mm}^2$ | Foramen Magnum |
| Uhasai et al. 2023<sup>14</sup> | $282.74 \pm 119.24 \text{ mm}^2$<br>*(Minor ellipse axis: $a = 7.50 \pm 2.50 \text{ mm}$)*<br>*(Major ellipse axis: $b = 12.0 \pm 3.10 \text{ mm}$)* | Fourth Ventricle |

Piaggio et al. 2018 observed a cross sectional area of 88.9 ± 6.0 mm^2^ at the foramen magnum^17^, which was used to convert the Yildiz et al. velocities to peak CSF flows during resting state of 1.45 ± 0.594 mL/s and during deep breathing of 2.04 ± 0.794 mL/s^1^ at the foramen magnum. We can then assume that this flowrate at the foramen magnum or 1^st^ cervical layer is equivalent to the flowrate at the fourth ventricle. This allows the computation of fourth ventricle fluid velocity by dividing each of these flowrates by the cross-sectional area of the fourth ventricle, as described by Uhasai et al. 2023^14^ as 282.74 ± 119.24 mm^2^. Thus, the Baselli et al. study estimates 0.393 ± 0.502 mm/s mean CSF flow caudally and the Yildiz et al. study estimates mean CSF flow velocity during resting state as 0.518 ± 0.646 mm/s and during breath holding as 0.918 ± 0.850 mm/s. Standard deviations were propagated across all steps using standard propagation of uncertainty for multiplicative equations.

## Notes

### Competing Interest Statement

The authors have declared no competing interest.

### Summary of Updates

Methods section on Experimental flow study and fMRI was changed to exclude the nonlinear fit to patient data in substitution for including a more robust nonlinear model of velocity reconstruction from literature. Figure 3B was altered for graphical consistency of the True Velocity. Methods section on In vivo data processing now includes an expanded description of the nonlinear model. Figure 4 was changed to demonstrate the differences between the linear, non-linear, and true velocity profiles. Figure 5 was changed to reflect the updated nonlinear model. Appendix sections were added to describe the nonlinear modeling framework and cerebrospinal fluid flow velocity comparisons.

